# The acidic domain of the hepatitis C virus NS4A protein is required for viral assembly and envelopment through interactions with the viral E1 glycoprotein

**DOI:** 10.1101/350132

**Authors:** Allison E. Roder, Stacy M. Horner

## Abstract

Hepatitis C virus (HCV) assembly and envelopment are coordinated by a complex protein interaction network that includes most of the viral structural and nonstructural proteins. While the nonstructural protein 4A (NS4A) is known to be important for viral particle production, the specific function of NS4A in this process is not well understood. We performed mutagenesis of the C-terminal acidic domain of NS4A and found that mutation of several of these amino acids prevented the formation of the viral envelope, and therefore the production of infectious virions, without affecting viral RNA replication. In an overexpression system, we found that NS4A interacted with several viral proteins known to coordinate envelopment, including the viral E1 glycoprotein. One of the NS4A C-terminal mutations, Y45F, disrupted the interaction of NS4A with E1. Specifically, NS4A interacted with the first hydrophobic region of E1, a region previously described as regulating viral particle production. Supernatants from HCV NS4A Y45F transfected cells had significantly reduced levels of HCV RNA, however they contained equivalent levels of Core protein. Interestingly, the Core protein secreted from these cells formed high order oligomers with a density matching the infectious virus secreted from WT cells. These results suggest that this Y45F mutation in NS4A causes secretion of low density Core particles devoid of genomic HCV RNA. These results corroborate previous findings showing that mutation of the first hydrophobic region of E1 also causes secretion of Core complexes lacking RNA, and therefore suggest that the interaction between NS4A and E1 is involved in the incorporation of viral RNA into infectious HCV particles. Our findings define a new role for NS4A in the HCV lifecycle and help elucidate the protein interactions necessary for production of infectious virus.

**Author Summary:** RNA viruses, which encompass both established and emerging pathogens, pose significant public health challenges. Viruses in the family *Flavivirdae*, including Dengue virus, Zika virus and hepatitis C virus (HCV), continue to cause morbidity and mortality worldwide. One HCV protein, NS4A, has known functions in several steps of the viral lifecycle, however, how it contributes to viral particle production is not understood. Here, we investigated the role of one region of NS4A, the C-terminal acidic domain, in regulating the viral lifecycle. We found that some of the amino acids within this domain are important for viral envelopment to make infectious particles, specifically through interaction with the E1 glycoprotein. NS4A interacts with the first hydrophobic domain of E1. Disruption of this interaction prevents the production of infectious virus particles and instead results in release of low density Core protein complexes that lack HCV RNA into the cellular supernatant. Overall, our results reveal that NS4A is important for late stages of the HCV lifecycle and suggest that the interaction between NS4A and E1 may regulate the incorporation of viral RNA into the virion for the formation of infectious HCV particles.

## Introduction

Hepatitis C virus (HCV) is a positive-sense RNA virus of the genus *Hepacivirus* in the *Flaviviridae* family. Over 70 million people worldwide are chronically infected with HCV and this chronic infection can lead to liver cirrhosis and hepatocellular cancer [1]. In the years spanning 2003-2013, HCV-related deaths numbered more than any other CDC-reported infectious disease [2]. Despite the availability of newly designed, highly effective direct-acting antivirals, disease prevalence remains high, and no vaccine exists for the virus [3–5].

HCV encodes a single stranded, positive-sense RNA genome of approximately 9.6 kilobases in length. Upon virus entry into hepatocytes, the viral genome is translated to form a single polyprotein. The polyprotein is co- and post-translationally cleaved by both host and viral proteases, including the NS3-NS4A viral protein complex, to form ten individual proteins. These ten proteins include both structural proteins, which eventually make up the virion, and non-structural proteins, which coordinate RNA replication and the other steps in the viral lifecycle, including virion assembly and envelopment (reviewed in [6]).

The late stages of the viral lifecycle, including assembly and envelopment, are just beginning to be dissected. While many details of these processes are not understood, recent work has uncovered several key steps that lead to production of infectious virus. Following RNA replication, HCV RNA is shuttled to the cytosolic lipid droplet where Core protein accumulates, oligomerizes, and recruits the NS3 and NS5A proteins [7–10]. NS5A is thought to play a role in RNA recruitment to the lipid droplet, whereas NS3 binds to Core and likely aids in movement of Core bound to RNA from the lipid droplet to nearby sites on the endoplasmic reticulum (ER) [11–16]. This process is coordinated by the NS2 protein which acts as a bridge between the non-structural protein NS3, and the structural protein E2, to link virion assembly at the lipid droplet to envelopment at the ER [17–22]. The role of NS2 in these steps is supported by the actions of the p7 protein [23, 24]. Lastly, Core oligomers bound to RNA bud into the ER lumen, acquiring an ER-derived lipid bilayer envelope that contains the viral E1 and E2 transmembrane glycoproteins [25, 26]. It is unclear what signals are necessary for the membrane curvature that results in budding but it is clear that E1 and E2 are necessary for successful envelopment, as deletion of E1 and E2 prevents formation of the viral envelope and production of infectious virions [24]. Following virion budding into the ER lumen, the virion is transported through the very-low-density lipoprotein (VLDL) secretory pathway acquiring apolipoproteins and other lipids, and is ultimately released from the cell in a noncytolytic manner as a lipoviroparticle [27–30]. In addition to the viral proteins mentioned here, a number of host proteins also play a role in HCV morphogenesis [31](reviewed in [32]).

In addition to its roles in viral assembly and envelopment, the NS3-NS4A protein complex has several other well-established functions in the HCV lifecycle. It is essential for viral polyprotein processing, viral RNA replication, and negative regulation of antiviral innate immunity (reviewed in [33]). NS3 functions as both a serine protease and an RNA helicase and requires its cofactor NS4A to enhance these activities and to target the protease complex to intracellular membranes [34, 35]. NS4A is 54 amino acids long and contains three domains, an N-terminal transmembrane domain that anchors NS3 to intracellular membranes, a central NS3-interaction domain required for proper folding of NS3, and a C-terminal domain that contains a kink region followed by an acidic region with a high number of acidic amino acids [36–39]. While the specific roles of NS4A in the HCV lifecycle are largely thought to occur indirectly through its function as a cofactor for NS3, mutation-based studies of the NS4A acidic domain suggest some independent roles for NS4A in regulating the HCV lifecycle, including during assembly and envelopment, as described below [36, 40].

There is strong evidence that NS3 and NS4A are each involved in the steps of virion assembly and envelopment. Specifically, NS3 has been shown to be involved in viral particle production through interactions with both Core and NS2 [15, 21, 22, 41]. In addition, culture adaptive amino acid mutations in the a0 helix of NS3 have been shown to promote viral assembly [16]. Separately, HCV particle production is also regulated by specific amino acids in the NS4A acidic domain such that when mutated, infectious virus formation is inhibited without affecting RNA replication. Some of these mutations can be partially rescued by compensatory amino acid substitutions in NS3, suggesting that NS3 and NS4A together can cooperate to regulate HCV particle production [40]. However, both the extent of the role of NS4A in assembly and envelopment and the specific function of NS4A in regulating the production of infectious HCV remain unclear.

Here, we define a new role for the NS4A protein in regulating HCV envelopment. We have identified amino acids in the acidic domain of NS4A that are required for the formation of the viral envelope. Further, we have found that NS4A alone can interact with a number of viral proteins that coordinate viral envelopment, including Core, E1, E2 and NS5A. Interestingly, disruption of the NS4A-E1 interaction prevents envelopment of the HCV particle and results in secretion of Core particles that are not associated with viral RNA. Taken together, our findings reveal a new role for NS4A in coordinating the HCV lifecycle and define new viral interactions that lead to successful HCV particle envelopment.

## Results

### A Y45F mutation in the hepatitis C virus NS4A protein causes a decrease in infectious viral titer

The acidic domain of NS4A has significant sequence homology between all HCV genotypes, with amino acids 40-54 in the acidic domain differing by at most 3 amino acids (Fig 1A). In particular, the tyrosine residue at position 45 is conserved in all seven genotypes of HCV. While previous studies have implicated the acidic domain of NS4A in regulating HCV RNA replication and particle production, the mechanism of this regulation was not explored [40]. We sought to investigate how the NS4A acidic domain contributes to HCV particle production. We engineered a structurally conservative amino acid substitution, changing the tyrosine residue (Y, TAT) at position 45 to a phenylalanine (F, TTT) in the genotype 2a strain of HCV, Japanese fulminant hepatitis-1 (JFH1) [42]. We then generated WT or NS4A Y45F *in vitro* transcribed RNA and transfected it into Huh7.5 cells. At 3 days post-transfection, while the WT RNA produced more than 3 logs of infectious virus, no titer was detected from cells transfected with the RNA containing the Y45F mutation, as measured by focus forming assay (Fig 1B). After several passages, Y45F RNA began to produce infectious virus, and after 14 days it produced equivalent titers to that of the WT virus (Fig 1B). Sequencing of the NS4A region of HCV RNA extracted from cells at 1, 3, and 14 days post-transfection revealed that the Y45F mutation had reverted back to WT by day 14, with some reversion detected as early as day 3 (Fig 1C). These results reveal that substitution of the tyrosine at position 45 with phenylalanine in NS4A prevents production of infectious HCV, indicating that Tyr-45 is required for the production of infectious virus.

**Fig 1.**
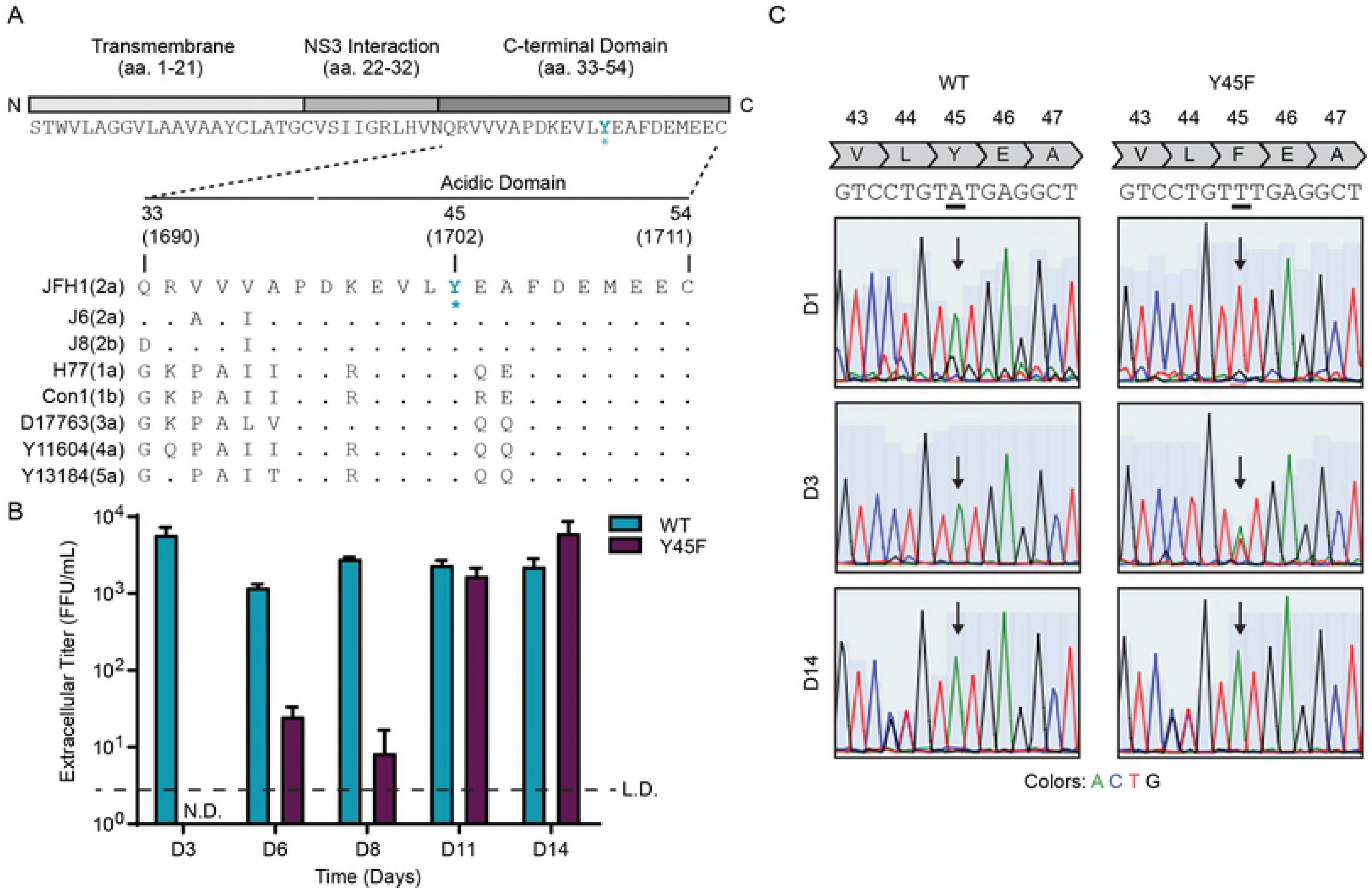
A Y45F mutation in hepatitis C virus NS4A causes a decrease in infectious viral titer. (A) Schematic of the viral NS4A protein. * indicates location of Y45F mutation. Numbers correspond with aa position within the NS4A protein (aa1-54) or the full-length polyprotein (aa1690-1711) (B) Focus forming assay of supernatants harvested from Huh7.5 cells at indicated days post electroporation with HCV WT or NS4A Y45F *in vitro* transcribed RNA. FFU/mL = focus forming units/milliliter. Values are presented as mean ± SD (n = 3). Data is representative of three independent experiments. (C) Sequencing of the NS4A region (nt 4710-5251) of cDNA amplified from Huh7.5 cells transfected with HCV WT or NS4A Y45F RNA in (B) at indicated days.

### The NS4A Y45F mutation in HCV does not alter viral RNA replication

To determine if the loss of infectious HCV production by the NS4A Y45F amino acid change was due to altered HCV RNA replication, we engineered the Y45F mutation into an HCV subgenomic replicon construct containing a luciferase reporter and measured luciferase production over time following transfection of Huh7.5 cells with *in vitro* transcribed HCV RNA. We found that the HCV replicon RNA with the Y45F mutation in NS4A replicated as efficiently as WT, while the HCV RNA with a lethal mutation in the NS5B RNA dependent RNA polymerase (GND) did not replicate (Fig 2A). Additionally, the HCV proteins NS3, NS4A and NS5A were expressed in lysates harvested at 48 hours post-transfection of either WT or Y45F RNA, indicating that the Y45F mutation did not affect the production of these viral proteins (Fig 2B). Of note, the epitope of the NS4A antibody is in the C-terminal domain of NS4A that contains Tyr-45. Therefore, the lack of a detectable NS4A band by immunoblotting in the mutant condition suggests that the Y45F mutation prevents NS4A recognition by this antibody (Fig 2B). Indeed, the fact that HCV RNA replication is not altered by the Y45F mutation indicates that the NS4A protein must be stably expressed, as NS4A is required for HCV RNA replication [36, 40, 43]. Since the interaction of NS3 with NS4A is essential for viral replication, we tested if the Y45F mutation impacted this interaction using a co-immunoprecipitation experiment with overexpressed proteins. The results show that the Y45F mutation does not alter NS3-NS4A complex formation (Fig 2C). Together, these results indicate that the Y45F mutation in NS4A does not alter HCV RNA replication, HCV protein expression, or NS3-NS4A complex formation. Therefore, the NS4A Y45F mutation in HCV must cause a defect at a later stage of the viral lifecycle.

**Fig 2.**
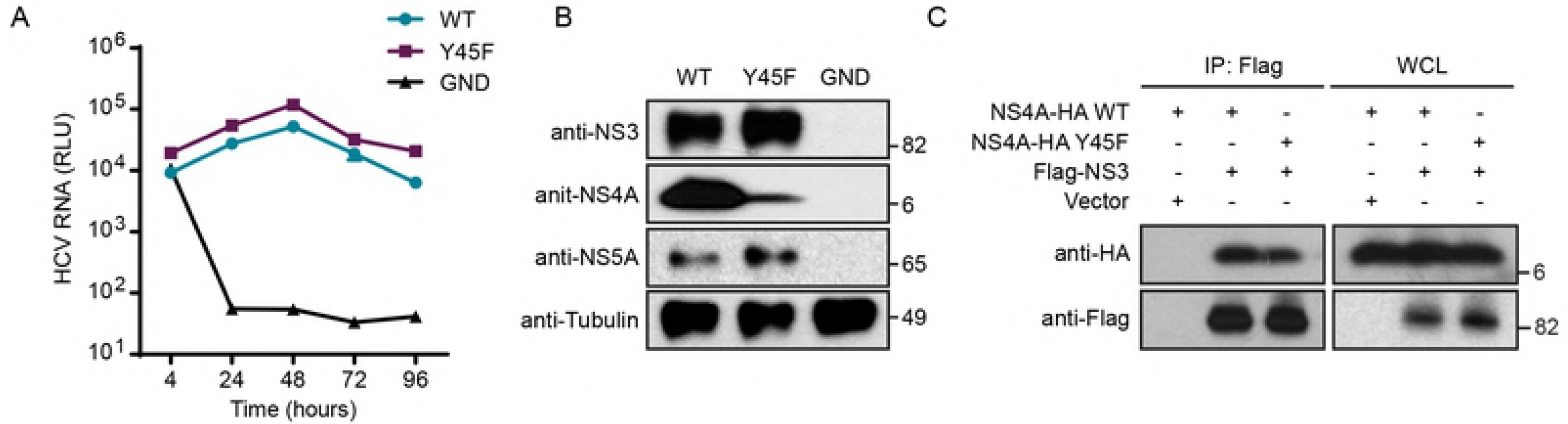
The NS4A Y45F mutation in HCV does not alter viral RNA replication. (A) Renilla luciferase assay to measure HCV replicon luciferase reporter (JFH1-SGR-luc) activity from Huh7.5 cells following electroporation. GND = lethal mutation in HCV NS5B RNA-dependent RNA polymerase. RLU = *Renilla* luciferase units. Data is presented as mean ± SEM, n=3. (B) Immunoblot analysis of extracts of Huh7.5 cells at 72 hours post transfection with indicated HCV RNA. (C) Immunoblot analysis of anti-Flag immunoprecipitated extracts and whole cell lysates (WCL) from Huh7.5 cells transfected with indicated tagged HCV proteins or vector.

### The NS4A Y45F mutation inhibits viral envelopment

As the NS4A Y45F mutation did not alter HCV RNA replication but did prevent infectious virus production, we next tested if this mutation affected viral assembly and envelopment or viral release. We first examined if the Y45F mutation caused a viral release defect by measuring both intracellular and extracellular titer. We transfected Huh7.5 cells with WT, Y45F, or GND HCV RNA, and measured the viral titer from the supernatant (extracellular titer) or from lysates generated by freeze-thaw cycles (intracellular titer) by using a focus forming assay. As before, HCV NS4A Y45F RNA did not produce extracellular titer (Fig 3A), and here we found that it also did not produce intracellular titer (Fig 3B). Taken together, these results indicate that the Y45F mutation impairs viral particle production prior to the formation of fully infectious virions.

**Fig 3.**
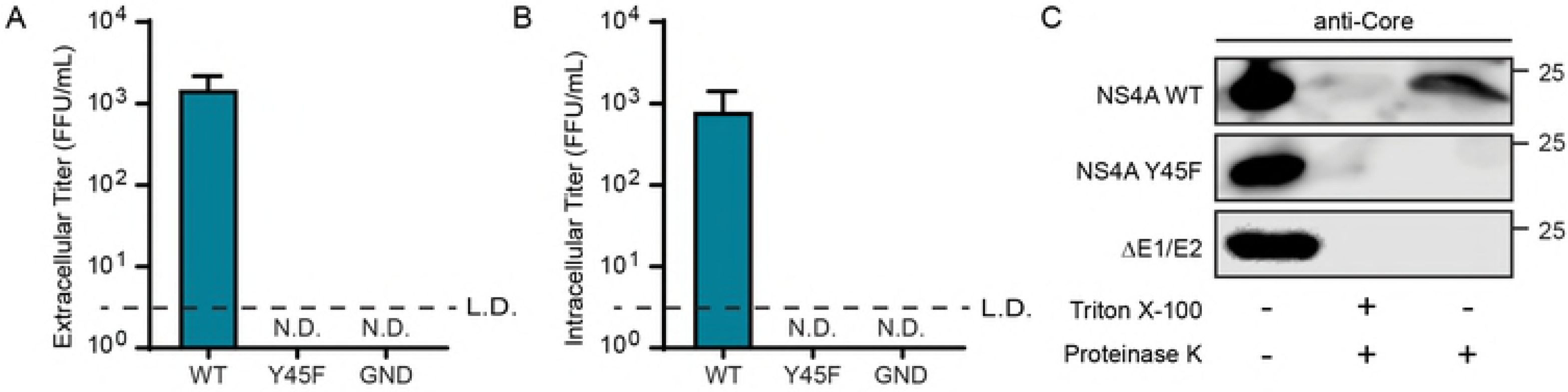
The NS4A Y45F mutation inhibits HCV envelopment. (A, B) Focus forming assay of supernatants for extracellular titer, or cellular lysates for intracellular titer, respectively, from Huh7.5 cells at 48 hours post electroporation with HCV WT, Y45F, or GND *in vitro* transcribed RNA. (C) Immunoblot analysis for HCV Core protein of cell lysates of Huh7.5 cells at 48 hours post electroporation with *in vitro* transcribed HCV RNA subjected to the indicated treatments in a proteinase K protection assay. ΔE1/E2 = complete deletion of E1 and E2 coding regions (aa 192 to 720 of JFH1). For panels A and B, data is presented as mean ± SEM, (n=3). C is representative of 3 independent experiments.

An infectious HCV virion contains viral RNA, encapsidated by the viral Core protein, surrounded by an outer lipid envelope containing the viral glycoproteins, E1 and E2, and cellular lipids and lipoproteins [32, 44, 45]. Taking advantage of these structural properties of the HCV virion, we next tested if the Y45F mutation prevented viral envelopment by using a proteinase K protection assay. The HCV Core protein in enveloped virions, which have an outer lipid envelope, is protected from degradation following proteinase K treatment [24]. Because the HCV glycoproteins are required for acquisition of the lipid bilayer membrane, a viral RNA with a deletion of the E1 and E2 coding region (amino acids (aa) 192-720, ΔE1/E2) can be used a negative control for envelopment [24]. Lysates were harvested from HCV RNA (WT, Y45F, or ΔE1/E2) transfected Huh7.5 cells, incubated with proteinase K, and analyzed by immunoblot for Core. We found that while Core was protected from proteinase K digestion in WT, it was not protected in lysates containing the Y45F mutation, similar to ΔE1/E2 (Fig 3C). These data indicate that the Y45F mutation prevents envelopment of the virion, resulting in a lack of both intracellular and extracellular viral titer, suggesting that Tyr-45 may be an important residue for HCV envelopment.

### The acidic domain of NS4A is required for HCV envelopment

Based on our findings that HCV RNA with the NS4A Y45F mutation has a defect in viral envelopment, we hypothesized that other amino acids in the NS4A C-terminal acidic domain may also be required. To test this, we introduced several mutations into the acidic domain of NS4A that were previously found to be important for production of infectious HCV particles (K41A, L44A, and E52A) and tested their effects on viral envelopment [40]. We performed a proteinase K protection assay, as in Figure 3, and found that the K41A, L44A and E52A mutants all resulted in a quantifiable decrease in protease-resistant Core as compared to WT, suggesting that these mutations also caused a defect in envelope formation (Figs 4A and 4B).

**Fig 4.**
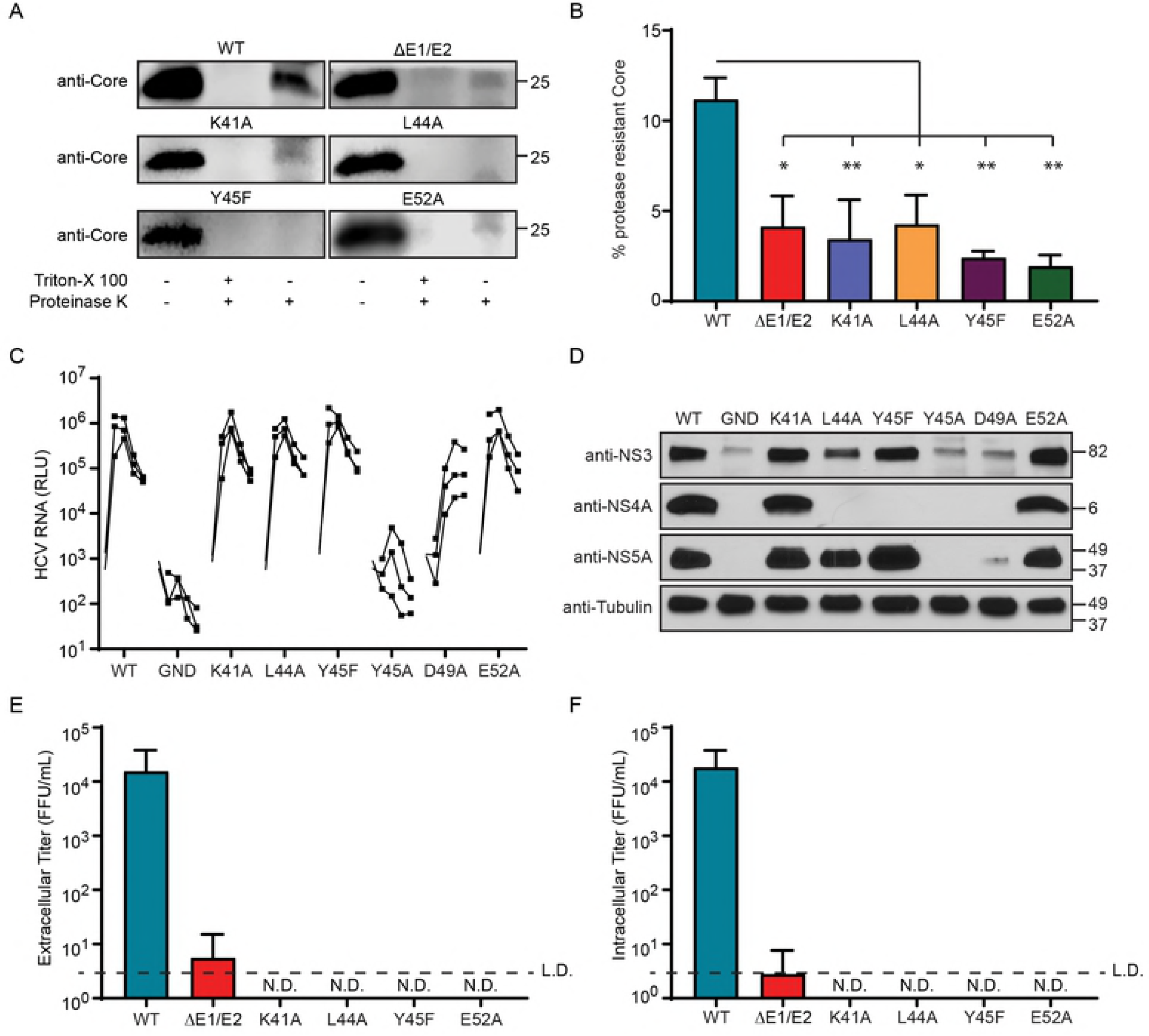
The acidic domain of NS4A is important for HCV envelopment. (A) Immunoblot analysis for HCV Core protein of cell lysates of Huh7.5 cells at 48 hours post electroporation with *in vitro* transcribed HCV RNA containing the indicated mutations in the NS4A region, subjected to the indicated treatments in a proteinase K protection assay. (B) Quantification of ratio of protease resistant Core to untreated Core from (A). ΔE1/E2 = complete deletion of E1 and E2 coding regions (aa 192 to 720 of JFH1). Data is presented as mean ± SEM (n=3). * - P < 0.05. ** - P < 0.01. (C) *Renilla* luciferase assay to measure HCV replicon luciferase reporter (JFH1-SGR-luc) activity from Huh7.5 cells following electroporation of indicated constructs. GND = lethal mutation in HCV NS5B RNA-dependent RNA polymerase. RLU = *Renilla* luciferase units. (D) Immunoblot analysis of lysates from 72 hour time point from (C). (E, F) Focus forming assay of supernatants for extracellular titer or cellular lysates for intracellular titer, respectively, from Huh7.5 cells electroporated *in vitro* transcribed HCV RNA containing the indicated mutations in the NS4A region. For (B, C, E, F), data is presented as mean ± SEM, (n=3). Panel A is representative of three independent experiments.

We additionally tested the impact of these amino acids on RNA replication, HCV protein expression, and production of both intracellular and extracellular titer; and also tested two additional mutations with known replication defects, Y45A and D49A, as controls [40]. HCV RNA containing the NS4A K41A, L44A, and E52A mutations all replicated and expressed HCV proteins to a similar extent as WT, while NS4A D49A and Y45A showed mild to severe replication defects (Figs 4C and 4D). These mutations all prevented intracellular and extracellular infectious virus from being produced, as seen previously by others (Figs 4E and 4F) [40]. Taken together, these data show that multiple amino acids within the acidic domain of NS4A are important for formation of the viral envelope and production of infectious virus.

### NS4A Y45 is required for NS4A interaction with the E1 glycoprotein

Because a complex network of HCV proteins regulates HCV assembly and envelopment, we hypothesized that NS4A may facilitate an interaction between either structural (Core, E1 or E2) or non-structural (p7, NS2 or NS5A) proteins to regulate these processes. Therefore, we first tested if overexpressed NS4A WT or Y45F interacted with Core, E1, or E2 using co-immunoprecipitation in Huh7.5 cells. We found that overexpressed NS4A WT interacts with Core, E1 and E2 (Figs 5A, S1A-B). While Core and E2 interactions with NS4A were equivalent for WT and Y45F (Figs S1A-B), the NS4A Y45F mutation greatly decreased NS4A and E1 interaction (Figs 5A-B). To determine if NS4A WT interacts with E1 also in the context of HCV infection we transfected Huh7.5 cells with an infectious clone of HCV containing an N-terminal HA tag on E1 [41]. We then immunoprecipitated E1 using the HA epitope and found that, indeed, NS4A and E1 can interact during HCV infection (Fig 5C). We also tested the interactions of NS4A with p7, NS2, and NS5A, non-structural proteins that all have roles in HCV envelopment [10–12, 17–21, 23, 24]. NS4A did interact with NS5A and this interaction was not altered by the Y45F mutation in NS4A (Fig. S1D). We found no interaction between overexpressed NS4A WT or Y45F with either NS2 or p7 (Fig S1C). Together, these data show that NS4A can bind to Core, E1, E2, and NS5A, and that mutation of NS4A at Tyr-45 disrupts its binding to the E1 protein.

**Fig 5.**
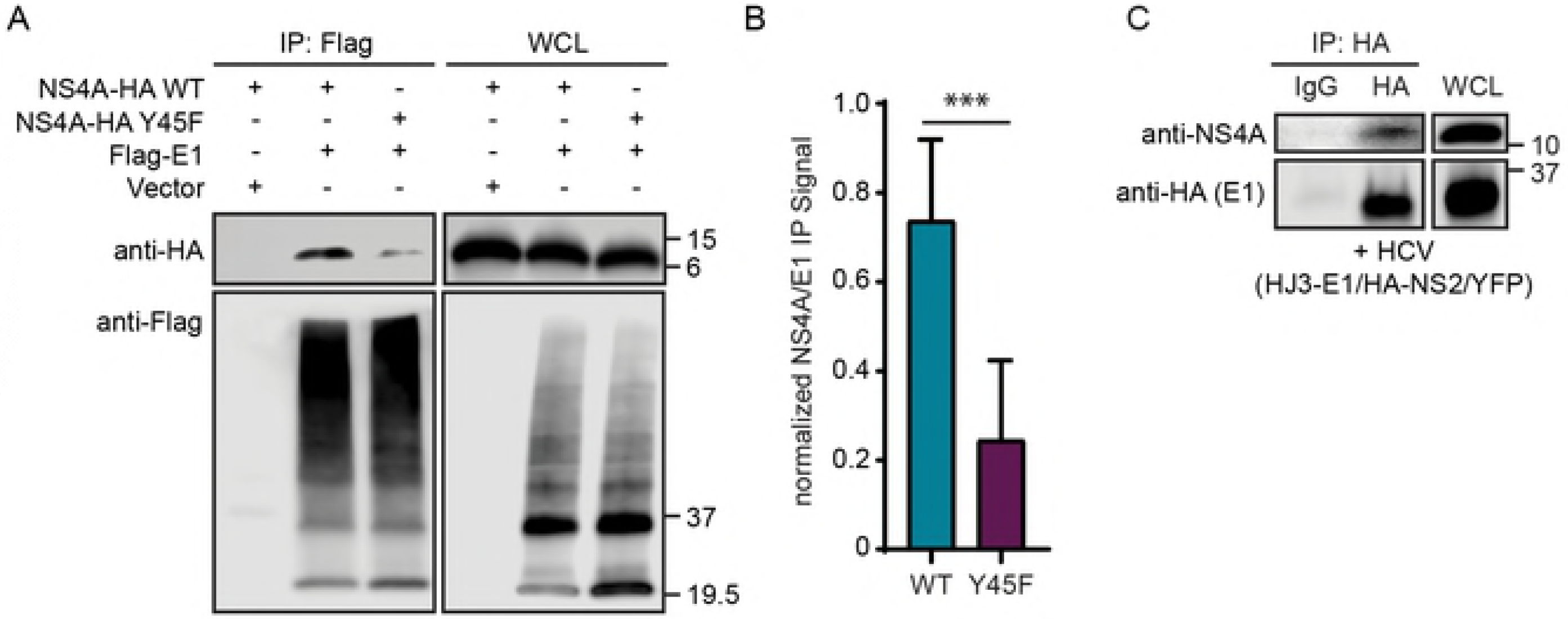
NS4A Y45 is required for NS4A interaction with the E1 glycoprotein. (A) Immunoblot analysis of anti-Flag immunoprecipitated extracts from Huh7.5 cells transfected with NS4A-HA WT, NS4A-HA Y45F and Flag-tagged E1. (B) Quantification of the NS4A:E1 signal seen in the immunoprecipitation from (A). Data is presented as mean ± SEM, (n=3). *** - P < 0.001. (C) Immunoblot analysis of anti-HA immunoprecipitated extracts from Huh7.5 cells at 48 hours post electroporation with *in vitro* transcribed HCV HJ3-E1/HA-NS2/YFP [41] which contains an N-terminal HA tag on the E1 protein.

### NS4A binds to the first hydrophobic region of E1

To investigate the mechanism of how the NS4A-E1 interaction might facilitate viral envelopment, we mapped the binding site of NS4A on E1. The E1 and E2 glycoproteins are translated in the ER membrane and are cleaved from the viral polyprotein by a host protease. After cleavage, E1 and E2 form a stable heterodimer and are retained in the ER. E1 has an N-terminal ectodomain, two hydrophobic regions and a C-terminal transmembrane domain (Fig 6A) (reviewed in [46]). We therefore created a series of E1 truncation mutants based on these known domains of E1, containing N-terminal Flag tags (Fig 6A). We overexpressed the truncation mutants and NS4A-HA in Huh7.5 cells and then performed Flag immunoprecipitations followed by immunoblotting for NS4A-HA. We found that NS4A co-immunoprecipitated with E1 aa1-106, aa1-138, and aa67-192, all of which contain the first hydrophobic region of E1, but did not interact with aa1-66 or aa107-192, which lack this region (Fig 6B). These data suggest that NS4A binds to E1 via the first hydrophobic region of the protein.

**Fig 6.**
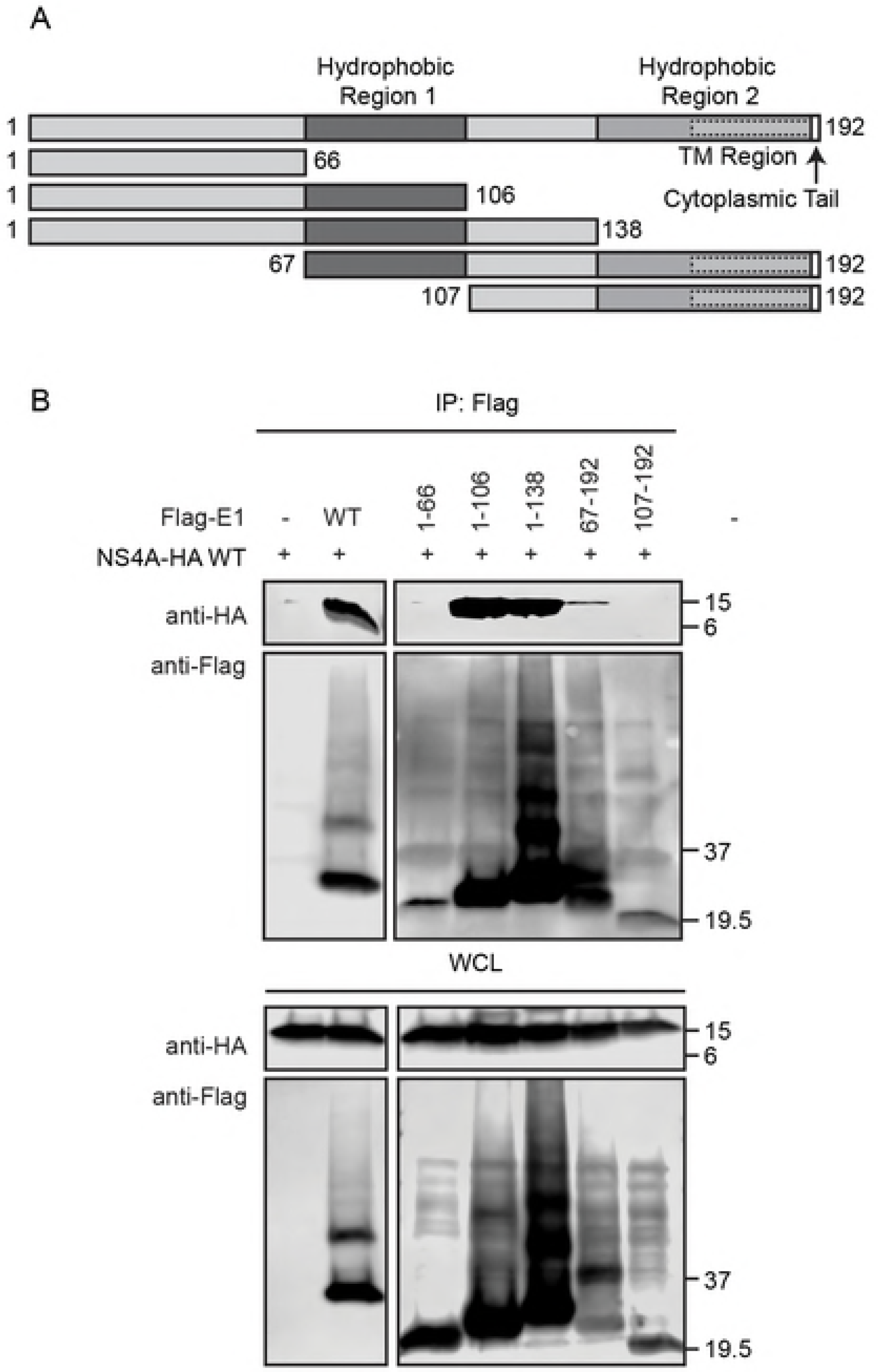
NS4A binds the first hydrophobic region of E1. (A) Schematic of the HCV E1 protein with functional domains indicated. aa locations are based on the JFH1 genome, within the E1 coding region. (B) Immunoblot analysis of anti-Flag immunoprecipitated extracts from Huh7.5 cells transfected with NS4A-HA WT and Flag-tagged E1, either full length (WT) or as indicated.

### NS4A Y45F results in release of Core oligomers devoid of HCV RNA

The first hydrophobic region of E1 has previously been implicated in viral particle production, and several amino acids within this region are important for viral infectivity [47, 48]. Additionally, one specific E1 mutation, D263A (E1 aa71), attenuated viral infectivity and resulted in secreted Core protein but not HCV RNA [48]. Taken together with our findings that NS4A binds to the E1 the hydrophobic region, which contains aaD263, we hypothesized that Y45F may have a similar phenotype as the E1 D263A mutant. Therefore, we measured the secretion of HCV RNA and Core protein into supernatants from cells replicating HCV NS4A WT or Y45F RNA. We found that while the Y45F mutation resulted in lower levels of extracellular HCV RNA, as measured by RT-qPCR, secretion of Core into the supernatant was unaltered as compared to WT (Figs 7A-B). These results were surprising as we did not expect to detect Core protein in the supernatant secreted from HCV NS4A Y45F transfected cells. Because of these unexpected data, we sought to profile the viral components in supernatants from Y45F cells. We collected and concentrated cellular supernatants from HCV NS4A WT or Y45F transfected cells and then ultracentrifuged these samples over iodixanol gradients. We collected 10 equal fractions from the top, with fraction 1 having the lowest density and fraction 10 having the highest density. Each fraction was analyzed by RT-qPCR for HCV RNA, by focus forming assay for viral titer, and by immunoblot for Core protein. In the WT samples, fractions 2 and 3 had the highest levels of both HCV RNA and viral infectivity (Fig 7C). These fractions also contained high molecular weight complexes of Core protein (Fig 7D, lanes 2 and 3). We observed a second peak of HCV RNA in fractions 7, 8, and 9, along with a small amount of higher density Core protein, however these fractions had significantly less viral infectivity (Fig 7D, lanes 7-9). Therefore, the HCV RNA in these higher density fractions is likely non-infectious and may represent secreted membrane-associated RNA from replication complexes [49]. However, we saw little to no infectious viral titer from any fraction in the Y45F samples and found the majority of HCV RNA present in fractions 7-9, while the expected infectious fractions (2-4) had little RNA (Fig 7C). Interestingly, high molecular weight complexes of Core protein were still observed in fractions 2-4, similar to the distribution of Core in WT samples (Fig 7D). The fact that Core forms oligomers and that these were in a different fraction than the peak of HCV RNA suggests that the Y45F mutation results in release of partially formed virions containing Core protein oligomers but devoid of HCV RNA.

**Fig 7.**
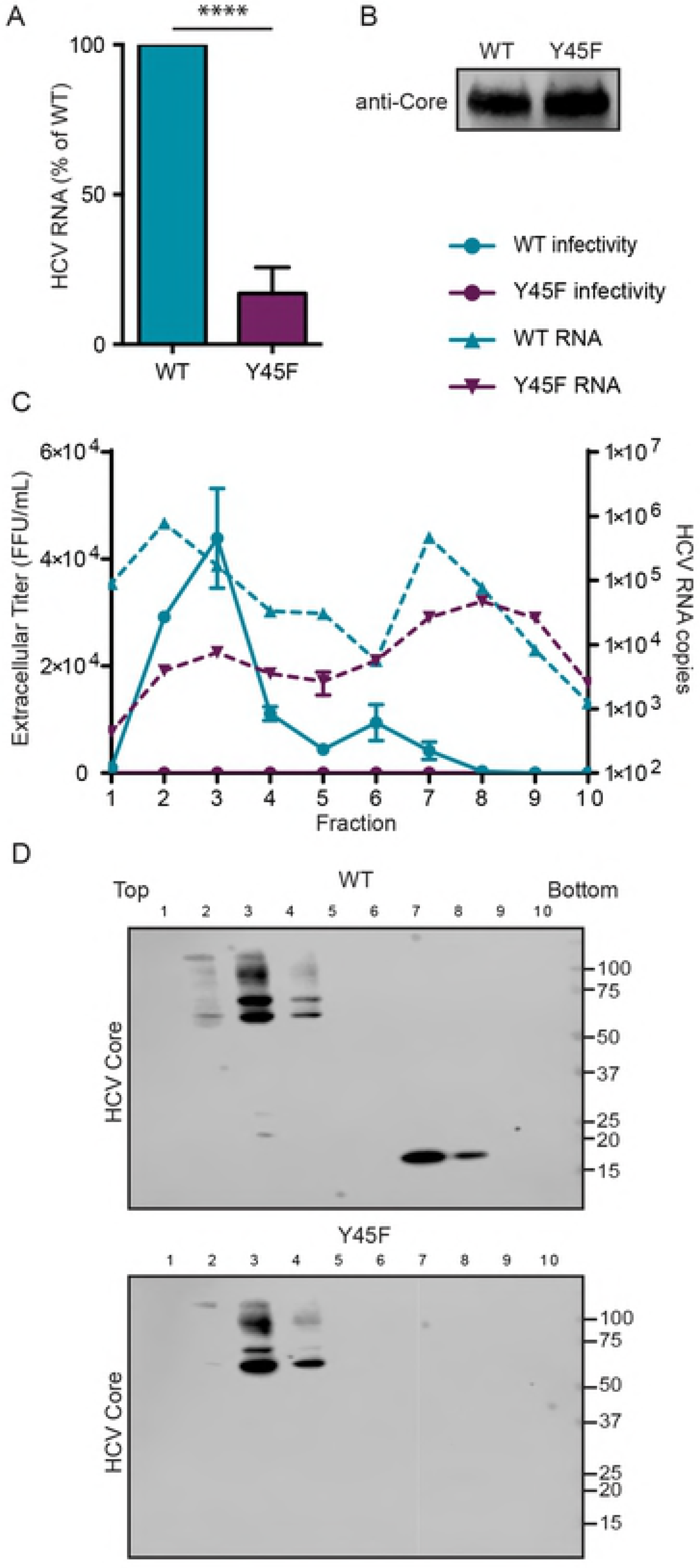
NS4A Y45F causes release of Core oligomers devoid of HCV RNA. Huh7.5 cells were electroporated with either WT or NS4A Y45F *in vitro* transcribed HCV RNA. Supernatants were harvested and analyzed for HCV RNA by RT-qPCR (A) or Core protein by immunoblot (B). Data in A are presented as mean ± SEM (n=4). **** - P < 0.0001. Supernatant from Huh7.5 cells electroporated with *in vitro* transcribed HCV WT or NS4A Y45F RNA was concentrated and fractionated over a 10-50% iodixanol gradient and 10 equal fractions were collected. Fractions were analyzed by focus-forming assay for infectivity and RT-qPCR for HCV RNA (C) and also for HCV Core protein by immunoblot (D). Fraction 1-10 correspond with fractions running from top to bottom of the gradient. B-D are representative of 2 independent experiments.

## Discussion

Our results define a new role for NS4A in the late stages of the HCV lifecycle. Specifically, we have found that the acidic domain of NS4A is important for regulating assembly and that mutation of specific amino acids within this domain prevents formation of the viral envelope. Further, we have identified new interactions between NS4A and both structural and non-structural viral proteins. This suggests that NS4A may act as a bridge, linking virion assembly at the lipid droplet and envelopment at the ER, similar to the actions of the NS2 protein. Importantly, we found that NS4A binds to E1 and that antagonizing this interaction with one amino acid change in NS4A prevents viral envelopment. We mapped the binding site of NS4A on E1 and found that it interacts with the first hydrophobic region, a region that is known to be important for viral particle production [47, 48]. Finally, we found that the Y45F mutation in NS4A, which prevents envelopment, also results in secretion of noninfectious, incompletely formed virions that are composed of low density Core protein oligomers that lack HCV RNA. Together our results reveal a new role for NS4A in coordinating the proper assembly and envelopment of HCV particles to make infectious virus.

The NS4A protein contains only 54 amino acids and yet has three distinct domains with specific functions in the HCV lifecycle. While the functions of the NS4A transmembrane domain and the NS3-interaction domain are largely defined [33], much less is known about the function of the C-terminal region, which contains a high number of acidic amino acids from aa40 to aa54 (Fig 1A). We found that mutation of several amino acids in the acidic domain, including Y45F, disrupted the formation of the viral envelope and therefore prevented production of infectious virus, without affecting viral RNA replication (Figs 2A, 3A, 3C, and 4). Indeed, the presence of a Tyr at aa45 of NS4A was so essential for the viral lifecycle that a viral RNA containing the Y45F mutation reverted back to the WT sequence after only a few days of passage in cell culture (Fig 1B-C). Given that changing a Tyr to Phe is a structurally conservative mutation, removing only the hydroxyl group, it is formally possible that NS4A could be phosphorylated at this tyrosine. However, it is unlikely that lack of phosphorylation of this tyrosine would be the sole contributor to the envelopment defects, as several other amino acids within the region displayed the same phenotypes when mutated (Fig 4) and a Y45A mutation prevented HCV RNA replication altogether (Fig 4C) [40]. Therefore, the acidic domain likely regulates envelopment through the concerted actions of the amino acids in this acidic region of NS4A.

Changing the amino acids in NS4A at K41, L44, Y45, and E52 to alanine all resulted in loss of viral titer due to defects in envelopment (Fig 4). The acidic domain of NS4A, which has been proposed to have an alpha helical structure, is important for replication, and indeed the Y45A change results in less replication [36, 40]. However, because mutation of the other amino acids within this C-terminal domain did not alter viral RNA replication, it is unlikely that they disrupt the conformation of the alpha helix. Structural predictions of this alpha helix suggest that K41, Y45 and E52 all lie on one face of the helix while L44 and D49 would face the opposite direction [36]. Therefore, it is possible that the amino acids we studied in this region could facilitate different protein interactions on opposite faces of the protein, each contributing to HCV envelopment. In support of this hypothesis, previous work has shown that an adaptive mutation in NS3 partially rescues an assembly defect resulting from the K41A mutation, suggesting that NS4A can cooperate with NS3 via K41 for viral particle production [40]. We found that NS4A WT and Y45F bound NS3 equivalently (Fig 2C) but that the Y45F mutation prevented NS4A interaction with E1 (Fig 5A). This suggests that while both the K41A and Y45F RNAs are defective in HCV envelopment, they may function in NS4A to facilitate different protein-protein interactions that regulate envelopment.

Supporting the hypothesis that NS4A interacts with several HCV proteins to coordinate virion envelopment, we did identify several previously unknown interactions of NS4A with both structural and non-structural proteins including Core, E1, E2, and NS5A. Others have shown that compensatory mutations within NS4A rescue assembly and envelopment defects caused by mutations in NS2, however we found that NS4A did not interact with NS2 during overexpression. Therefore, these data suggest that during infection, NS2 and NS4A likely work together through a multi-protein complex or to perform similar roles in the lifecycle [19]. Indeed, NS2 is considered to be the main organizer of envelopment, binding both structural and nonstructural proteins to link viral assembly steps at the lipid droplet to envelopment steps at the ER [17–21, 23, 24]. NS4A also binds to proteins involved in both early (Core, NS3, and NS5A) and later (E1 and E2) steps of assembly and envelopment which could suggest that NS4A may also serve as a link between virion production steps at the lipid droplet and the ER, similar to NS2. Overall, these results suggest that NS2 and NS4A could play similar roles in organizing and facilitating viral envelopment.

We found that NS4A binds to E1 and that this interaction is disrupted by the Y45F mutation (Fig 5), suggesting that the NS4A-E1 interaction is important for envelopment of the virion. The E1 protein has an N-terminal ectodomain, two internal hydrophobic domains, a transmembrane domain and a very short, 2 amino acid cytoplasmic, C-terminal tail (Fig 6A) [46]. Surprisingly, we found that NS4A binds to the first hydrophobic region of the protein (Fig 6B). This region has also previously been shown to bind to Core, and mutations within this domain diminish viral particle production [47, 48, 50–52]. Curiously, this E1 domain is hydrophobic and is proposed to associate with the lipid membrane bilayer while the NS4A acidic domain is cytoplasmic and not known to have membrane interactions. It is possible that the acidic domain of NS4A could associate with the ER membrane to interact with this region of E1. However, it is equally likely that NS4A and E1 are linked by a host protein. Future studies designed to determine how NS4A interacts with E1 would yield further insights into the HCV envelopment process.

The finding that NS4A binds to the first hydrophobic region of E1 is particularly interesting, as this region in E1 has recently been shown to regulate viral particle production [47, 48]. In fact, a D263A mutation at the start of hydrophobic region 1 in E1 (E1 aa71), resulted in decreased viral titer and secretion of Core particles that were devoid of genomic RNA. Further, this mutation disrupted the localization of E1 with HCV RNA in fluorescence *in situ* hybridization experiments [48]. In our studies, fractionation of supernatant from cells replicating HCV NS4A Y45F RNA revealed that low density fractions contained little to no HCV RNA, similar to E1 D263A (Fig. 7). However, these low density fractions contained secreted Core oligomers, suggesting that these oligomers were associated with cellular lipids or apolipoproteins. Indeed, transfected Core protein has been shown to self-assemble into higher order complexes and non-enveloped particles have been found in the serum of HCV infected patients [53, 54]. Transfection of Core alone can also alter VLDL secretion, and therefore it is possible that secreted Core may be associated with cellular lipids and lipoproteins [55]. Taken together, these data suggest that the NS4A acidic domain and the E1 first hydrophobic domain cooperate during envelopment, perhaps to aid in the incorporation of viral RNA into the virion. While NS4A itself does not have RNA binding capability, it is linked to NS3, which contains an RNA binding helicase domain, and thus, NS3-NS4A together could cooperate via E1 interactions for incorporation of RNA into the developing virion.

Our study contributes new insights into the steps required for HCV to form infectious viral particles. As the viral particle lifecycle stages that occur in association with lipid droplets and the ER are tightly linked and likely occur nearly simultaneously, it is unclear if nucleocapsid intermediates (Core protein assembled around HCV RNA) exist separate from fully enveloped nucleocapsids [32]. Our data show that Core protein assembles into oligomers prior to envelopment and suggest that the function of NS4A in viral assembly and envelopment is after this Core oligomerization step. Further, the fact that we identified Core protein oligomers that did not contain a protective envelope or HCV RNA suggests that RNA incorporation into the virion at or near envelopment sites could be a necessary signal for virion budding events to occur. Our data therefore support a model by which NS4A interacts with E1 to link viral RNA to Core oligomers in the forming virion and signal the envelopment of the Core-RNA complex.

## Materials and Methods

### Cell lines and culture conditions

Huh7.5 cells, which have been previously described [56], were maintained in Dulbecco’s modification of eagle’s medium (DMEM; Mediatech) supplemented with 10% fetal bovine serum (FBS; HyClone) and 25mM HEPES (Thermo-Fisher) at 37°C with 5% CO_2_. The identity of the Huh7.5 cells used in this study was verified by using the Promega GenePrint STR kit (DNA Analysis Facility, Duke University), and cells were verified as mycoplasma free by the LookOut Mycoplasma PCR detection kit (Sigma).

### Plasmids and site-directed mutagenesis

These plasmids have been described previously: psJFH1-p7+NS [57] and HJ3-E1/HA-NS2/YFP ([41], gift of Dr. MinKyung Yi). psJFH1-p7+NS is a culture adapted strain of JFH1 containing 7 mutations within p7 and the nonstructural proteins [57]. pJFH1-SGR-luc contains a bicistronic replicon as follows: [JFH1-derived untranslated region (UTR; nt 1-397)]-[in frame *Renilla* luciferase reporter]-[EMCV IRES-nonstructural genes (NS3-NS5B)]. To make this plasmid, a DNA fragment encoding *Renilla* luciferase was fused between the T7 promoter sequence-5’ UTR of JFH1 and the EMCV IRES-nonstructural genes from pSGR-JFHI [58] following PCR (for oligonucleotide sequence see Table 1), digestion (with inserted *BglII* site between 5’UTR and 5’end of *Renilla*, a *PmeI* site between the 3’end of *Renilla* and 5’end of the ECMV IRES, and an existing *AgeI* site in the 5’UTR of pSGR-JFH1), and a 3-piece ligation. Mutagenesis of constructs was performed using the QuikChange lightning site-directed mutagenesis kit (Stratagene) on pJFH1-SGR-luc or psJFH1-p7+NS using the indicated oligonucleotides (Table 1). psJFH1-p7+NS ΔE1/E2 was constructed by removing amino acids 192-720 from the psJFH1 p7+NS background [24]. HCV over-expression constructs (noted below) were constructed by PCR amplification of the gene of interest from psJFH1-p7+NS and insertion of the *Pmel-Notl* digested fragment into pEF-Tak-Flag [59] or the *EcoRl-Xbal* digested fragment into pEF1. pEF-Tak Flag-NS2 was created using InFusion (Clontech) after PCR. Table 1 provides the sequence of all oligonucleotides used. Bold letters in the oligonucleotide sequences indicate overlap with vector sequence, and the sequence of the HA tag within the oligonucleotides is underlined. All nucleotide and amino acid positions refer to the JFH1 genome (GenBank accession number: AB047639). The sequences of all plasmids were verified by DNA sequencing and are available upon request.

**Table 1:**
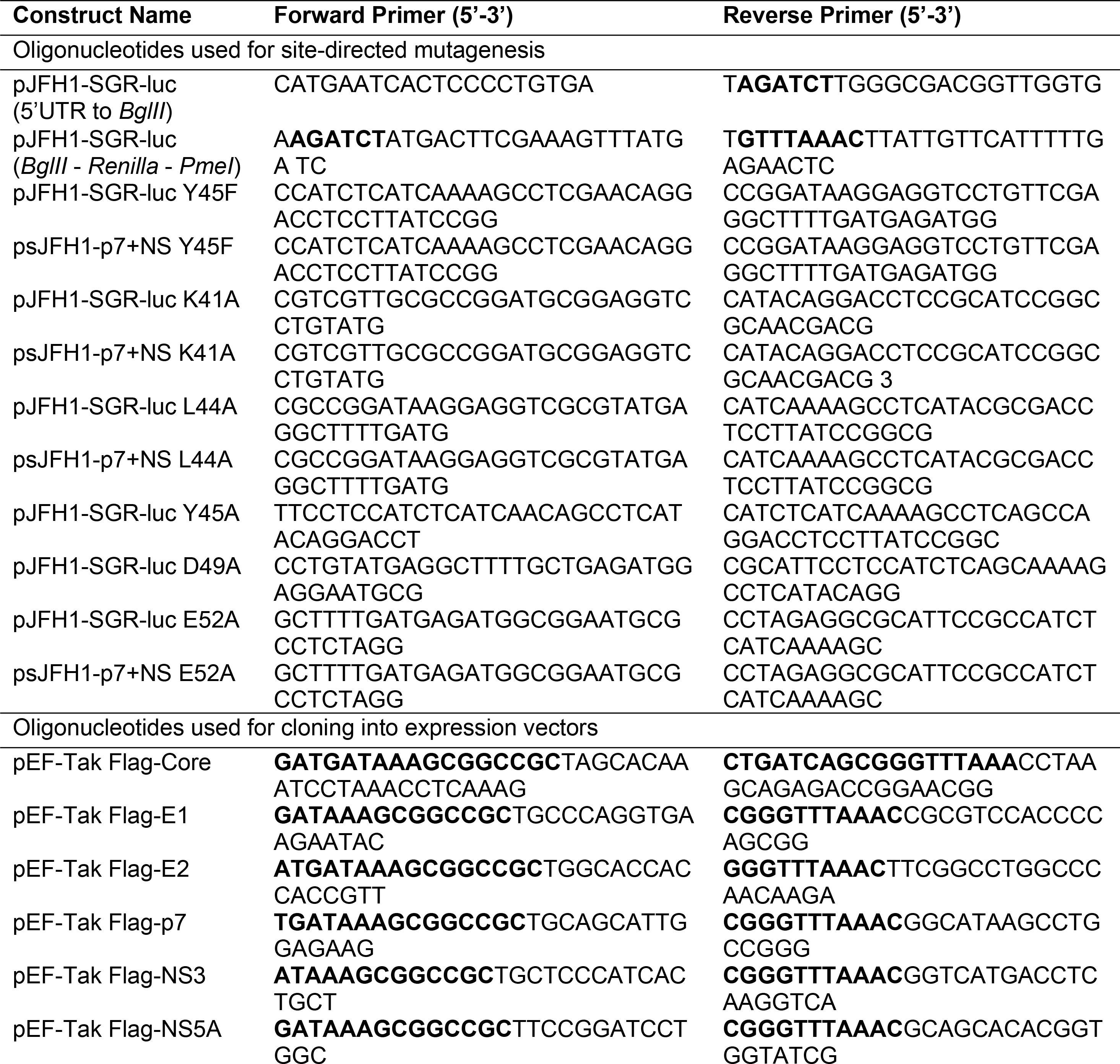
Oligonucleotides used in this study.

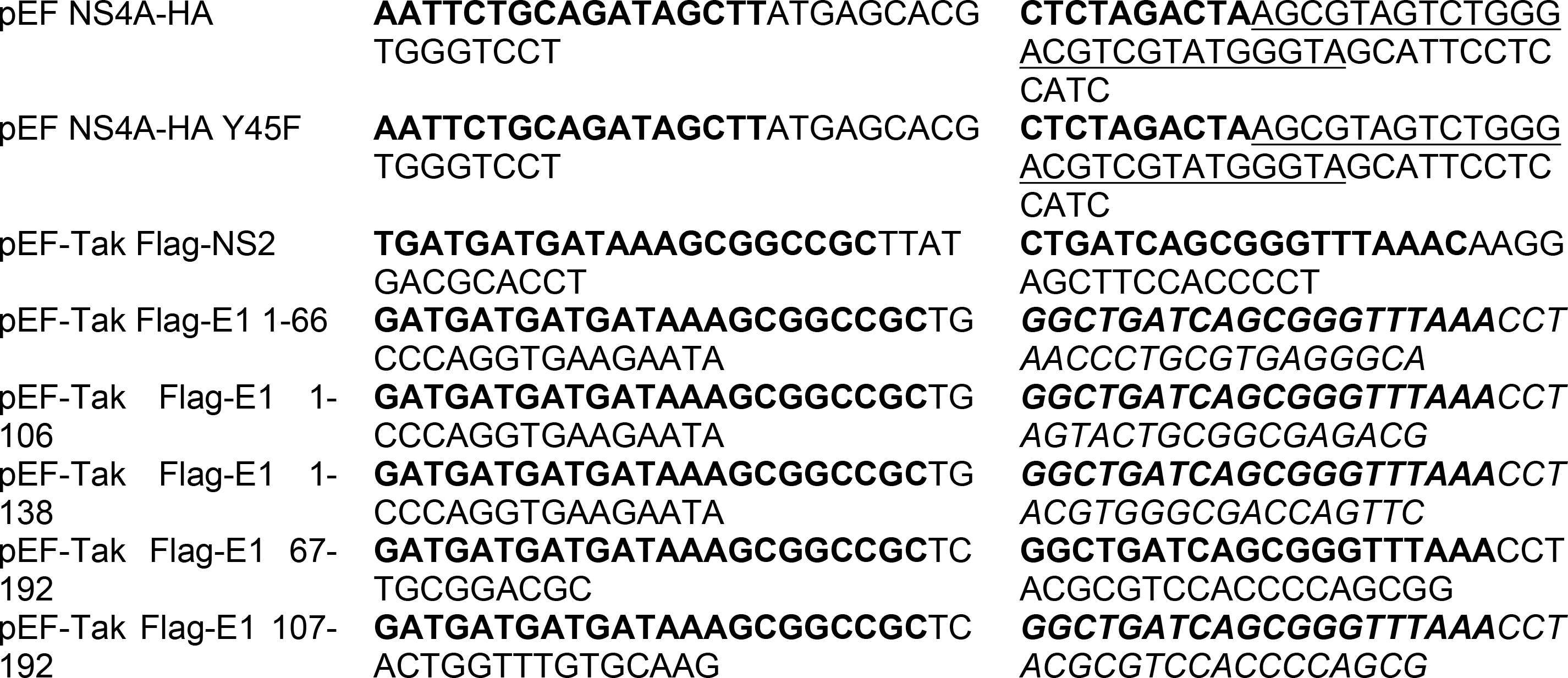

### *In vitro* transcription of HCV RNA and electroporation

Plasmid DNA encoding the described HCV constructs was linearized using the *Xbal* restriction enzyme. Purified linearized DNA was used as a template for *in vitro* transcription with a MEGAscript T7 transcription kit (Thermo-Fisher). The *in vitro* transcribed RNA was treated with DNase (Thermo-Fisher) and then purified by phenol-chloroform extraction. The quality of the RNA was verified on a denaturing gel. For electroporation, 1μg (HCV Replicon RNA) or 5μg (HCV Infectious Clone RNA) was electroporated into 4×10^6^ Huh7.5 cells in Cytomix electroporation buffer (120mM KCl, 10mM Potassium Phosphate Buffer, 5mM MgCl_2_, 25mM HEPES, 0.15mM CaCl_2_, 2mM EGTA, pH 7.6) at 250V and 950μF in a 4mm cuvette with a Gene Pulser Xcell system (Bio-Rad). Four hours post electroporation, cells were washed extensively with Phosphate Buffered Saline (PBS) and cDMEM.

### Focus forming assay

#### Extracellular titer

Supernatants were harvested from Huh7.5 cells electroporated with HCV RNA at indicated time points, serially diluted, and used to infect naïve Huh7.5 cells in triplicate wells of a 48-well plate for 2 hours. Plates were harvested at 48hpi and fixed with 4% paraformaldehyde. Cells were permeabilized (0.2% Triton-X-100 in PBS), blocked (10% FBS in PBS) and immunostained with mouse anti-HCV NS5A antibody (9e10, 1:500, gift of Dr. Charles Rice). Infected cells were visualized following incubation with horseradish peroxidase (HRP)-conjugated secondary antibody (1:500; Jackson ImmunoResearch) and VIP Peroxidase Substrate Kit (Vector Laboratories). Foci were counted at 40X magnification. Titer (FFU/mL) was determined as previously described [60]. *Intracellular titer.* Cell pellets were washed with PBS and resuspended in serum-free DMEM. Cells were then lysed using a series of freeze/thaw cycles in a dry ice/Ethanol bath. Post-nuclear supernatants were used to infect naïve Huh7.5 cells, and a focus forming assay was performed as described above.

### Luciferase assays

JFH1 SGR-luc *in vitro* transcribed RNA (1μg) was electroporated into Huh7.5 cells. Cells were suspended in 20mL cDMEM and plated in 12-well plates. Cells were harvested after a PBS wash by incubation in *Renilla* lysis buffer (Promega). *Renilla* luciferase values were measured according to manufacturer’s instructions (Renilla Luciferase Assay System, Promega) using a BioTek Synergy 2 microplate reader.

### HCV NS4A sequencing

RNA was extracted from cells by using the Qiagen RNeasy kit according to manufacturer’s instructions and then used as a template for cDNA synthesis with the iScript cDNA synthesis kit (BioRad). The NS4A region of the HCV genome was amplified by nested PCR with the following oligonucleotides. Round 1. 5’ – CAGTCCGATGGAGAAGAAGG - 3’, 5’ - GCATGGGATGGGGCAGTC - 3’, Round 2. 5’ - ACACATAGACGCCCACTTCC - 3’, 5’ - GTATGTCCTGGGCCTGCTTA - 3’, and then the 542bp PCR product was purified and sequenced by Sanger sequencing.

### Quantification of HCV RNA by RT-qPCR

RNA from cells was isolated using the RNeasy kit (Qiagen) and RNA from infected supernatants was isolated using the QIAamp viral RNA kit (Qiagen), both according to manufacturer’s instructions. The RNA copy number of harvested RNA was measured in triplicate by RT-qPCR using the TaqMan Fast Virus 1-Step Mix (Qiagen) with an HCV-specific probe targeting the 5’ untranslated region of HCV (Assay ID: Pa03453408_s1). The copy number was calculated by comparison to a standard curve of a full-length *in vitro* transcribed HCV RNA, as described [57].

### Immunoblotting

Cells were lysed in a modified radio immunoprecipitation assay (RIPA) buffer (10mM Tris pH 7.5, 150mM NaCl, 0.5% sodium deoxycholate, 1% Triton X-100) supplemented with protease inhibitor cocktail (Sigma) and phosphatase inhibitor cocktail (Millipore), and post-nuclear supernatants were harvested by centrifugation. Quantified protein was resolved by SDS/PAGE, transferred to PVDF membranes using the Turbo-transfer system (BioRad) and blocked with StartingBlock (Thermo-Fisher) or 3% bovine serum albumin (Sigma) in PBS with 0.1% Tween (PBS-T). Membranes were probed with specific antibodies, washed with PBS-T and incubated with species-specific HRP conjugated antibodies (Jackson ImmunoResearch, 1:5000), washed again with PBS-T, and treated with Pico PLUS enhanced chemiluminescent (ECL) reagent (Thermo-Fisher). The signal was then captured on X-ray film or by using a LICOR Odyssey FC. Antibodies used for immunoblot include mouse anti-HCV Core (1:250, Abcam), mouse anti-HCV NS3 (1:500, Abcam), rabbit anti-HCV NS4A (1:1000, Genscript [61]), mouse anti-HCV NS5A (1:500, 9e10, gift of Dr. Charles Rice), anti-Flag HRP (1:2500, Sigma), and rabbit anti-HA (1:500, Sigma).

### Proteolytic Protection Assay

This protocol was adapted from the manuscript by Gentzsch and colleagues [24]. Briefly, cells electroporated with JFH1-p7+NS *in vitro* transcribed RNA were harvested at 48 hours post electroporation by scraping into cold proteinase K buffer (50mM Tris-HCl pH 8.0, 10 mM CaCl_2_, 1mM DTT). Cells were then lysed by five freeze/thaw cycles and aliquots of lysate (50μL) were either (i) left untreated (ii) pretreated with 5μL of 10% Triton-X-100 followed by proteinase K treatment (50μg/mL) for 30 minutes on ice or (iii) treated with proteinase K only. Proteinase K treatment was terminated by incubation with 10mM phenylmethane sulfonyl fluoride. The samples were mixed with 4X SDS sample buffer (1M Tris (pH 6.8), 60% glycerol, 0.06% Bromophenol Blue, 12% SDS)), incubated at 50°C for 5 minutes, and immunoblotted for HCV Core protein, as above.

### Immunoprecipitations

300-500μg of protein extracted as above was incubated with 50μL anti-Flag M2 magnetic beads (Sigma) in 1X Tris buffered saline (TBS) at 4°C overnight with head over tail rotation. Beads were washed 3X in modified 1X RIPA buffer and eluted in 2X Laemmli Buffer (BioRad) at 50°C for 5 minutes. Protein was resolved by SDS/PAGE and immunoblotting, as above.

### Biochemical subcellular fractionation

Concentrated supernatants were purified over a 10-50% iodixanol gradient, as previously described [48]. Briefly, at 48 hours post electroporation of HCV RNA in Huh7.5 cells, supernatant was collected, mixed with polyethylene glycol (PEG) 8000 to a final concentration of 8% and incubated with rocking at 4°C overnight. PEG supernatants were centrifuged at 11,000 X g for 30 minutes, supernatant was removed, and remaining pellets were suspended in cold 1X PBS. These resuspensions were layered over a 10-50% iodixanol gradient and centrifuged at 222,000 X g in a SW41 rotor in a Beckman Coulter ultracentrifuge. 10 equal fractions (1ml) were collected with a BioComp piston gradient fractionator, and then viral titer (FFU/ml), HCV RNA copy number, and HCV Core protein (immunoblotting) was measured from each fraction.

### Statistical Analysis

Student’s unpaired t-tests and one-way analysis of variance (ANOVA) were used for statistical analysis of data. Values are presented as mean ± standard error of the mean (n=3 or as indicated). * - P < 0.05, ** - P < 0.01, *** - P < 0.001, **** - P < 0.0001.

## Acknowledgements

We thank all members of the Horner lab for helpful discussion; Christine Vazquez, Michael McFadden, Matthew Sacco, and Daltry Snider for reading of the manuscript; and Kevin Labagnara, Moonhee Park, Alexandra Fink and Mawuli Attipoe for assistance with experiments. We thank the following for reagents. Dr. Charles Rice at Rockefeller University and Dr. MinKyung Yi at University of Texas Medical Branch.

## Supporting information

**Fig S1. NS4A binds to Core and NS5A but not p7 or NS2.** Immunoblot analysis of anti-Flag immunoprecipitated extracts from Huh7.5 cells transfected with NS4A-HA WT, NS4A-HA Y45F and Flag-tagged Core (A), E2 (B), p7/NS2 (C), or NS5A (D).

## References

1. Global Hepatitis Report 2017. Geneva. World Health Organization; 2017. Licence. CC BY-NC-SA 3.0 IGO.

2. Ly KN, Hughes EM, Jiles RB, Holmberg SD. Rising Mortality Associated With Hepatitis C Virus in the United States, 2003-2013. Clin Infect Dis. 2016;62(10).1287–8.

3. Liang TJ, Ghany MG. Current and future therapies for hepatitis C virus infection. N Engl J Med. 2013;368(20).1907–17.

4. Mohd Hanafiah K, Groeger J, Flaxman AD, Wiersma ST. Global epidemiology of hepatitis C virus infection. new estimates of age-specific antibody to HCV seroprevalence. Hepatology. 2013;57(4).1333–42.

5. Li D, Huang Z, Zhong J. Hepatitis C virus vaccine development. old challenges and new opportunities. National Science Review. 2015;2(3).10.

6. Scheel TK, Rice CM. Understanding the hepatitis C virus life cycle paves the way for highly effective therapies. Nat Med. 2013;19(7).837–49.

7. Miyanari Y, Atsuzawa K, Usuda N, Watashi K, Hishiki T, Zayas M, et al. The lipid droplet is an important organelle for hepatitis C virus production. Nat Cell Biol. 2007;9(9).1089–97.

8. Matsumoto M, Hwang SB, Jeng KS, Zhu N, Lai MM. Homotypic interaction and multimerization of hepatitis C virus core protein. Virology. 1996;218(1).43–51.

9. Klein KC, Dellos SR, Lingappa JR. Identification of residues in the hepatitis C virus core protein that are critical for capsid assembly in a cell-free system. J Virol. 2005;79(11).6814–26.

10. Appel N, Zayas M, Miller S, Krijnse-Locker J, Schaller T, Friebe P, et al. Essential role of domain III of nonstructural protein 5A for hepatitis C virus infectious particle assembly. PLoS Pathog. 2008;4(3).e1000035.

11. Yin C, Goonawardane N, Stewart H, Harris M. A role for domain I of the hepatitis C virus NS5A protein in virus assembly. PLoS Pathog. 2018;14(1).e1006834.

12. Tellinghuisen TL, Foss KL, Treadaway J. Regulation of hepatitis C virion production via phosphorylation of the NS5A protein. PLoS Pathog. 2008;4(3):e1000032.

13. Han Q, Xu C, Wu C, Zhu W, Yang R, Chen X. Compensatory mutations in NS3 and NS5A proteins enhance the virus production capability of hepatitis C reporter virus. Virus Res. 2009;145(1):63–73.

14. Ma Y. NS3 helicase domains involved in infectious intracellular hepatitis C virus particle assembly. J Virol. 2008;82(15):7624–39.

15. Jones DM, Atoom AM, Zhang X, Kottilil S, Russell RS. A Genetic Interaction between the Core and NS3 Proteins of Hepatitis C Virus Is Essential for Production of Infectious Virus. J Virol. 2011;85(23):12351–61.

16. Yan Y, He Y, Boson B, Wang X, Cosset FL, Zhong J. A Point Mutation in the N-Terminal Amphipathic Helix alpha0 in NS3 Promotes Hepatitis C Virus Assembly by Altering Core Localization to the Endoplasmic Reticulum and Facilitating Virus Budding. J Virol. 2017;91(6).

17. Yi M, Ma Y, Yates J, Lemon SM. Compensatory mutations in E1, p7, NS2, and NS3 enhance yields of cell culture-infectious intergenotypic chimeric hepatitis C virus. J Virol. 2007;81(2):629–38.

18. Jones CT, Murray CL, Eastman DK, Tassello J, Rice CM. Hepatitis C virus p7 and NS2 proteins are essential for production of infectious virus. J Virol. 2007;81(16):8374–83.

19. Phan T. Hepatitis C virus NS2 protein contributes to virus particle assembly via opposing epistatic interactions with the E1-E2 glycoprotein and NS3-NS4A enzyme complexes. J Virol. 2009;83(17):8379–95.

20. Jirasko V, Montserret R, Lee JY, Gouttenoire J, Moradpour D, Penin F, et al. Structural and functional studies of nonstructural protein 2 of the hepatitis C virus reveal its key role as organizer of virion assembly. PLoS Pathog. 2010;6(12):e1001233.

21. Popescu CI, Callens N, Trinel D, Roingeard P, Moradpour D, Descamps V, et al. NS2 protein of hepatitis C virus interacts with structural and non-structural proteins towards virus assembly. PLoS Pathog. 2011;7(2):e1001278.

22. Stapleford KA, Lindenbach BD. Hepatitis C virus NS2 coordinates virus particle assembly through physical interactions with the E1-E2 glycoprotein and NS3-NS4A enzyme complexes. J Virol. 2011;85(4):1706–17.

23. Boson B, Granio O, Bartenschlager R, Cosset FL. A concerted action of hepatitis C virus p7 and nonstructural protein 2 regulates core localization at the endoplasmic reticulum and virus assembly. PLoS Pathog. 2011;7(7):e1002144.

24. Gentzsch J, Brohm C, Steinmann E, Friesland M, Menzel N, Vieyres G, et al. hepatitis c Virus p7 is critical for capsid assembly and envelopment. PLoS Pathog. 2013;9(5):e1003355.

25. Ai LS, Lee YW, Chen SS. Characterization of hepatitis C virus core protein multimerization and membrane envelopment: revelation of a cascade of core-membrane interactions. J Virol. 2009;83(19):9923–39.

26. Dubuisson J, Hsu HH, Cheung RC, Greenberg HB, Russell DG, Rice CM. Formation and intracellular localization of hepatitis C virus envelope glycoprotein complexes expressed by recombinant vaccinia and Sindbis viruses. J Virol. 1994;68(10):6147–60.

27. Chang T-HH. Human Apolipoprotein E Is Required for Infectivity and Production of Hepatitis C Virus in Cell Culture. J Virol. 2007;81(24):13783–93.

28. Jiang J, Luo G. Apolipoprotein E but not B is required for the formation of infectious hepatitis C virus particles. J Virol. 2009;83(24):12680–91.

29. Gastaminza P, Cheng G, Wieland S, Zhong J, Liao W, Chisari FV. Cellular determinants of hepatitis C virus assembly, maturation, degradation, and secretion. J Virol. 2008;82(5):2120–9.

30. Huang H, Sun F, Owen DM, Li W, Chen Y, Gale M, Jr., et al. Hepatitis C virus production by human hepatocytes dependent on assembly and secretion of very low-density lipoproteins. Proc Natl Acad Sci U S A. 2007;104(14).5848–53.

31. Coller KE, Heaton NS, Berger KL, Cooper JD, Saunders JL, Randall G. Molecular determinants and dynamics of hepatitis C virus secretion. PLoS Pathog. 2012;8(1).e1002466.

32. Bartenschlager R, Penin F, Lohmann V, Andre P. Assembly of infectious hepatitis C virus particles. Trends Microbiol. 2011;19(2).95–103.

33. Morikawa K, Lange CM, Gouttenoire J, Meylan E, Brass V, Penin F, et al. Nonstructural protein 3-4A. the Swiss army knife of hepatitis C virus. J Viral Hepat. 2011;18(5).305–15.

34. Tanji Y, Hijikata M, Satoh S, Kaneko T, Shimotohno K. Hepatitis C virus-encoded nonstructural protein NS4A has versatile functions in viral protein processing. J Virol. 1995;69(3).1575–81.

35. Failla C, Tomei L, De Francesco R. An amino-terminal domain of the hepatitis C virus NS3 protease is essential for interaction with NS4A. J Virol. 1995;69(3).1769–77.

36. Lindenbach BD, Pragai BM, Montserret R, Beran RK, Pyle AM, Penin F, et al. The C terminus of hepatitis C virus NS4A encodes an electrostatic switch that regulates NS5A hyperphosphorylation and viral replication. J Virol. 2007;81(17).8905–18.

37. Kim JL, Morgenstern KA, Lin C, Fox T, Dwyer MD, Landro JA, et al. Crystal structure of the hepatitis C virus NS3 protease domain complexed with a synthetic NS4A cofactor peptide. Cell. 1996;87(2).343–55.

38. Wolk B, Sansonno D, Krausslich HG, Dammacco F, Rice CM, Blum HE, et al. Subcellular localization, stability, and trans-cleavage competence of the hepatitis C virus NS3-NS4A complex expressed in tetracycline-regulated cell lines. J Virol. 2000;74(5).2293–304.

39. Brass V, Berke JM, Montserret R, Blum HE, Penin F, Moradpour D. Structural determinants for membrane association and dynamic organization of the hepatitis C virus NS3-4A complex. Proc Natl Acad Sci U S A. 2008;105(38).14545–50.

40. Phan T, Kohlway A, Dimberu P, Pyle AM, Lindenbach BD. The acidic domain of hepatitis C virus NS4A contributes to RNA replication and virus particle assembly. J Virol. 2011;85(3):1193–204.

41. Ma Y, Anantpadma M, Timpe JM, Shanmugam S, Singh SM, Lemon SM, et al. Hepatitis C virus NS2 protein serves as a scaffold for virus assembly by interacting with both structural and nonstructural proteins. J Virol. 2011;85(1):86–97.

42. Kato T, Furusaka A, Miyamoto M, Date T, Yasui K, Hiramoto J, et al. Sequence analysis of hepatitis C virus isolated from a fulminant hepatitis patient. J Med Virol. 2001;64(3):334–9.

43. Asabe SI, Tanji Y, Satoh S, Kaneko T, Kimura K, Shimotohno K. The N-terminal region of hepatitis C virus-encoded NS5A is important for NS4A-dependent phosphorylation. J Virol. 1997;71(1):790–6.

44. Bartosch B, Dubuisson J, Cosset FL. Infectious hepatitis C virus pseudo-particles containing functional E1-E2 envelope protein complexes. J Exp Med. 2003;197(5):633–42.

45. Nielsen SU, Bassendine MF, Burt AD, Bevitt DJ, Toms GL. Characterization of the genome and structural proteins of hepatitis C virus resolved from infected human liver. J Gen Virol. 2004;85(Pt 6):1497–507.

46. Op De Beeck A, Dubuisson J. Topology of hepatitis C virus envelope glycoproteins. Rev Med Virol. 2003;13(4):233–41.

47. Tong Y, Chi X, Yang W, Zhong J. Functional Analysis of Hepatitis C Virus (HCV) Envelope Protein E1 Using a trans-Complementation System Reveals a Dual Role of a Putative Fusion Peptide of E1 in both HCV Entry and Morphogenesis. J Virol. 2017;91(7).

48. Haddad JG, Rouille Y, Hanoulle X, Descamps V, Hamze M, Dabboussi F, et al. Identification of Novel Functions for Hepatitis C Virus Envelope Glycoprotein E1 in Virus Entry and Assembly. J Virol. 2017;91(8).

49. Pietschmann T, Lohmann V, Kaul A, Krieger N, Rinck G, Rutter G, et al. Persistent and transient replication of full-length hepatitis C virus genomes in cell culture. J Virol. 2002;76(8):4008–21.

50. Ma HC, Ke CH, Hsieh TY, Lo SY. The first hydrophobic domain of the hepatitis C virus E1 protein is important for interaction with the capsid protein. J Gen Virol. 2002;83(Pt 12):3085–92.

51. Lo SY, Selby MJ, Ou JH. Interaction between hepatitis C virus core protein and E1 envelope protein. J Virol. 1996;70(8):5177–82.

52. Nakai K, Okamoto T, Kimura-Someya T, Ishii K, Lim CK, Tani H, et al. Oligomerization of hepatitis C virus core protein is crucial for interaction with the cytoplasmic domain of E1 envelope protein. J Virol. 2006;80(22):11265–73.

53. Kunkel M, Lorinczi M, Rijnbrand R, Lemon SM, Watowich SJ. Self-assembly of nucleocapsid-like particles from recombinant hepatitis C virus core protein. J Virol. 2001;75(5):2119–29.

54. Maillard P, Krawczynski K, Nitkiewicz J, Bronnert C, Sidorkiewicz M, Gounon P, et al. Nonenveloped nucleocapsids of hepatitis C virus in the serum of infected patients. J Virol. 2001;75(17):8240–50.

55. Perlemuter G, Sabile A, Letteron P, Vona G, Topilco A, Chretien Y, et al. Hepatitis C virus core protein inhibits microsomal triglyceride transfer protein activity and very low density lipoprotein secretion: a model of viral-related steatosis. FASEB J. 2002;16(2):185–94.

56. Sumpter R, Jr., Loo YM, Foy E, Li K, Yoneyama M, Fujita T, et al. Regulating intracellular antiviral defense and permissiveness to hepatitis C virus RNA replication through a cellular RNA helicase, RIG-I. J Virol. 2005;79(5):2689–99.

57. Aligeti M, Roder A, Horner SM. Cooperation between the Hepatitis C Virus p7 and NS5B Proteins Enhances Virion Infectivity. J Virol. 2015;89(22):11523–33.

58. Kato T, Date T, Miyamoto M, Furusaka A, Tokushige K, Mizokami M, et al. Efficient replication of the genotype 2a hepatitis C virus subgenomic replicon. Gastroenterology. 2003;125(6).1808–17.

59. Saito T, Hirai R, Loo YM, Owen D, Johnson CL, Sinha SC, et al. Regulation of innate antiviral defenses through a shared repressor domain in RIG-I and LGP2. Proc Natl Acad Sci U S A. 2007;104(2).582–7.

60. Gastaminza P, Kapadia SB, Chisari FV. Differential biophysical properties of infectious intracellular and secreted hepatitis C virus particles. J Virol. 2006;80(22).11074–81.

61. Horner SM, Liu HM, Park HS, Briley J, Gale M Jr,. Mitochondrial-associated endoplasmic reticulum membranes (MAM) form innate immune synapses and are targeted by hepatitis C virus. Proc Natl Acad Sci U S A. 2011;108(35).14590–5.

